# Reduced leaf numbers due to shade avoidance allows sugar beet to maintain biomass under drought stress

**DOI:** 10.1101/2024.06.28.601301

**Authors:** Joe Ballenger, Albert T. Adjesiwor, Brent Ewers, Cynthia Weinig, Andrew R. Kniss

**Author notes:** Corresponding author’s.

## Abstract

The physical environment is complex even in managed agroecosystems, and plants often experience multiple stressors such as light competition and drought simultaneously. Phenotypic responses to multiple stressors may exhibit tradeoffs or constraints, such that harvestable yield is reduced In this study, we investigate how sugar beets respond to combinations of neighbor presence, resource competition, and water stress, and test the hypothesis that shade avoidance cues from neighboring vegetation may cause yield loss similar to, or even greater than, resource depletion by neighboring vegetation. Sugar beets were grown in 19L pails surrounded by Kentucky bluegrass (*Poa pratiensis*). Competition treatments included “shade avoidance only” where *B. vulgaris* leaves were exposed to reflected light from neighbors without shading or root competition, “shade avoidance plus root interaction” where crop and competitor roots interacted, and “no competition” as a control. Sugar beets in each competition treatment were subjected to 3 levels of irrigation: 100% of estimated evapotranspiration (ET), 80% ET and 60% ET. When grown without drought stress (100% ET), the shade avoidance treatment had 16% less root biomass compared to the no competition control (P = 0.007); however, under the most severe drought treatment (60% ET) root biomass was similar between the shade avoidance and no competition treatments (P = 0.48). Under 60% ET, plants in the shade avoidance only treatment produced 15% fewer leaves (P < 0.001) but 14% greater leaf area (P = 0.049) compared to the no competition control. The reduced number of leaves produced due to shade avoidance resulted in less structural tissue to be maintained under drought stress, and thus provides a potential benefit to ‘shade avoidance’ responses that are not directly related to avoiding shade.

## INTRODUCTION

Because plants are sessile organisms, phenotypic plasticity may significantly affect fitness in stressful environments. Given the complexity of the physical environment, plants must often initiate adaptive plastic responses to multiple types of stressors that they experience simultaneously. In arid and semi-arid regions, a combination of water stress and competition for light with other plants is common. Drought stress and shade avoidance result in opposing phenotypic responses. Drought stress elicits an investment in root tissue, which is needed to obtain water under the surface (Pierik & Testerink, 2014). Shade avoidance, on the other hand, results in an investment in shoot tissues at the expense of root development (Ruberti et al., 2012). Thus, understanding how these opposing morphological responses to neighbor sensing interact with water stress is integral to understanding the fitness consequences of plasticity to water-limited environments in settings where competition is also present.

The shade avoidance syndrome is a system by which plants perceive and acclimate to the presence of neighboring competitors. Plants sense close-proximity neighbors through changes in light quality, specifically the reduction of red (R) wavelengths and the enrichment of far-red (FR) wavelengths. Plants produce phytochrome (PHY) proteins that sense low R:FR and disable PIF transcription factors (Lorrain et al., 2008). Although there is a high degree of redundancy in phytochrome proteins, PHYB is primarily responsible for mediating shade avoidance responses (Casal, 2012; Franklin & Whitelam, 2004; Smith, 2000). Sensing a reduction in the ratio of R:FR light results in PHYB becoming inactive, leading to accumulation of PIF proteins and changes in transcriptional patterns (Lorrain et al., 2008). This change in transcriptional patterns results in a series of morphological adjustments referred to as the shade avoidance syndrome. These changes generally include reduced blade:petiole ratios, leaf hyponasty, decreased branching, greater internode elongation, and an investment in stem tissue at the expense of root growth (Bai et al., 2014; Casal, 2012; Green-Tracewicz et al., 2011; Schmitt et al., 2003).

The canonical shade avoidance response has been described in plants such as rice and soybean, crops selected for seed production which display tall growth patterns. These crops display classic internode elongation, which allows them to overtop neighbors (Britz, 1990; Christensen, 1995; Green-Tracewicz et al., 2011; Jannink et al., 2000; Lyu et al., 2021; Place et al., 2011; Wille et al., 2017). When exposed to far-red light indoors, agronomically important rosette-forming species lettuce and kale tend to grow wider in stance with reduced chlorophyll concentration in the leaves (Meng et al., 2019). Crops selected for root allocation, such as sugar beet, display reduced biomass, reduced leaf number, reduction in petiole and leaf length, and reductions in leaf area in response to shade avoidance (Adjesiwor et al., 2021; Adjesiwor & Kniss, 2020; Schambow et al., 2019).

In addition to neighbor sensing, plants also display morphological responses above and below ground to water stress. Plants move water from the soil column to the leaves through transpiration, requiring approximately 1000 times more water per carbon dioxide fixed in photosynthesis. Plants close guard cells to close stomata and limit transpiration to maintain sufficient leaf water potentials, but this reduces photosynthesis (Wong et al 1979). Osmotic adjustment, the accumulation of solutes from the soil or through synthesis to increase osmotic pressure, is another common response to water limitation (Turner, 2018). Absolute root growth is less under water limitations, but as long as there are sufficient resources plants will allocate relatively more to roots than shoots under drought, reflected in higher metabolism in roots than shoots under drought (Gargallo-Garriga et al., 2014). Plant roots can respond to differences in moisture content by growing towards areas of higher moisture, and reducing lateral root growth in areas of lower moisture (Bao et al., 2014; Dinneny, 2019).

Markesteijn and Poorter (2009) suggested that drought and shade tolerance are associated with different suites of traits, which are largely independent from one another. Specifically, dry forest species are thought to enhance their access to water in deeper soil layers by extending their root systems into these layers (Markesteijn & Poorter, 2009). Moist forest species are thought to enhance their light foraging capacity and increase nutrient acquisition through extending their leaves high up into the canopy or otherwise enhancing their light gathering capabilities (Markesteijn & Poorter, 2009). While the preceding morphological responses to shade and drought may be adaptive in trees, plants with other growth forms, such as herbaceous and particularly rhizomatous and rosette-forming species, will likely exhibit different morphological responses and potentially experience growth trade-offs. To the authors’ knowledge, no studies have addressed the potential tradeoffs in growth and yield between drought stress and shade avoidance of plants cultivated specifically for resource storage in the taproot.

Sugar beet, *Beta vulgaris,* is an important crop grown for its sucrose-rich taproot, and represents 20% of the world’s sugar supply. Drought stress, especially early in crop development, is an important determinant of final *B. vulgaris* root yield (Carter et al., 1980; Yonts et al., 2003). Season-long shade avoidance cues, even in the absence of water or nutrient deficiency, can cause between 30 to 70% biomass loss in sugar beet (Schambow et al., 2019). Similar to drought stress, early exposure to shade avoidance cues — especially between sugar beet emergence and the 2 true-leaf stage — are more impactful than late-season exposure to a similar environment (Adjesiwor et al., 2021). Understanding how the combination of drought stress and stress due to shade avoidance interact to affect growth will assist growers in formulating weed control and irrigation decisions.

Optimal Growth Theory (OPT) is the idea that plants adjust their growth patterns based on what resource is needed most. For example, studies in European beech (*Fagus sylvatica*) demonstrate that fine root biomass increases in response to reductions in precipitation (Hertel et al., 2013). The idea guiding the interpretation of the data in Hertel et al 2013 is because the needed resource is water, located underground, that the plants extended their root systems in search for this resource. Taking a similar view of the phenomenon of shade avoidance, the plants extend themselves further into the canopy via stem elongation because the needed resource is higher in the canopy. However, caution must be taken when interpreting results with certain crops, namely root crops, due to their strict developmental programs which preferentially shift resources to underground storage organs (Coleman & McConnaughay, 1995). This view of plant growth has recently been challenged by the Local Allocation Model (LAM) of plant growth (Robinson, 2023). The LAM suggests that growth rate does not reflect goal-seeking behavior, but rather reflects constraints imposed by resource availability. A reduced investment in fibrous root mass due to neighbor sensing could hypothetically decrease a plant’s ability to procure underground resources during competition under the OPT (Rajcan et al., 2004; van Gelderen et al., 2018). However, because plants allocate proportionally more to root systems under water stress, these effects could be offset depending on the amount of water stress and perhaps reflect the non-optimal allocation of resources to roots and leaves in line with the LAM (Padilla et al., 2009; Robinson, 2023).

Sugar beets are grown in the irrigated semi-arid regions of the Western US, a region marked by water scarcity, so understanding the interplay between drought stress and shade avoidance may impact irrigation, weed control, and cover crop management recommendations. While the negative impact of weeds are well understood in agriculture, the comparative importance of competition for belowground resources and developmental changes due to neighbor sensing are not well understood for crops which have been bred to supply resource-rich taproots. The objective of this experiment is to determine how plants respond differently when challenged with stressors whose responses result in conflicting growth patterns such as drought stress and shade avoidance. We hypothesized sugar beets would preferentially allocate growth to roots under drought stress and allocate growth to shoots upon the sensing of FR light reflected from neighbors.

## MATERIALS AND METHODS

### Experimental Setup

Studies were conducted outdoors during 2018 and 2019 at the University of Wyoming Laramie Research and Extension Center (LREC), Laramie, WY. The study comprised a factorial arrangement of 3 competition treatments (no competition, shade avoidance only, and shade avoidance plus root interaction), and 3 levels of irrigation (60, 80, 100% of estimated evaoptranspiration needs). The study was designed as a randomized complete block with 30 replicates of each competition by irrigation combination, and location in the field used as a blocking criteria. Black plastic 19 L pails (Figure 1) were filled with a potting mix (Berger, BM Custom Blend, Saint-Modeste, QC Canada) leaving about 7.5 cm head-space (Adjesiwor et al., 2021; Green-Tracewicz et al., 2011; Schambow et al., 2019). This soil was then fertilized with 85 g pail^-1^ of 14:14:14:5.5% (N:P:K:S) polymer-coated fertilizer (Florikote^TM^ NPK, Florikan E.S.A.-LLC, Sarasota, FL) to ensure adequate release of nutrients throughout the growing season and minimize nutrient limitations to plant growth.

**Figure 1.**
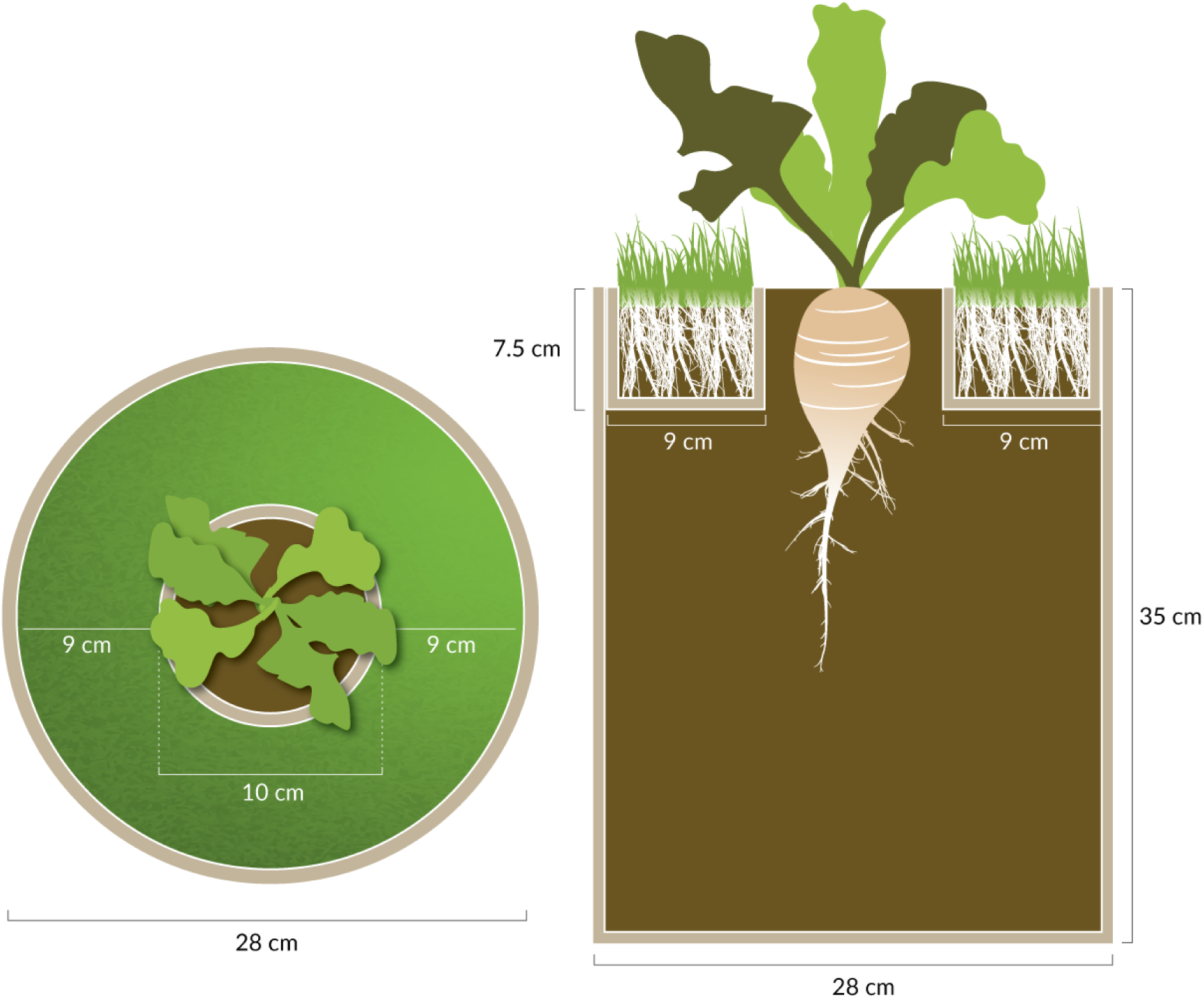
Illustration of the grass treatment used in the field experiment from the top-view (left) and a cross-section (right), modelled after (Green-Tracewicz et al., 2011), as modified by (Schambow et al. 2019). Sugar beet was planted into the center ring, which was created using a cardboard tube. Grass roots were constrained to the top 7.5 cm and outer 9 cm of the pail, and were isolated from the sugar beet roots using a plastic barrier. The soil treatment was designed identically, except no grass was planted into the potting media in the outer ring. Drawing by Jessica Perry.

A cardboard tube (10 cm diameter Staples Kraft Heavy-Duty Mailing Tube: Staples Inc., Framingham, MA) of height 7.5 cm was taped to a 2-mil thick white plastic bag and laid on top of the potting mix such that the cardboard tube was in the middle of the pail and top of the tube was flush with the rim of the pail (Figure 1). Kentucky bluegrass (*Poa pratensis* L.) sod was planted in the ring on top of the plastic for competition treatments, or filled with potting media for no-competition treatments. The plastic prevented roots of the grass from interacting with sugar beet roots in the shade avoidance only treatment. For the shade avoidance plus root interaction treatment, the plastic layer was perforated with 24 holes of 6.5mm diameter each, allowing grass roots to access the potting media and interact with the sugar beet root. Potting mix was then added into the cardboard tube to be flush with the top of the cardboard tube. Three sugar beet seeds (cultivar ‘W322NT’) were planted per pail in the center of the cardboard tube. Emerged sugar beet seedlings were thinned to one seedling per pail within 48 h after emergence. Grass was trimmed by hand at least a weekly basis or as otherwise needed to ensure sugar beet plants were not exposed to direct shade from the grass treatments.

### Irrigation

Sugar beet was drip irrigated from planting until harvest to ensure proper application of moisture treatments. All irrigation treatments for beets were watered using emitters which exuded 2 liters/hour. Grass in the shade avoidance only treatment was irrigated as needed using separate drip irrigation lines using 4 liters/hour emmitters. To determine average daily sugar beet water use, sugar beet were grown in 19 L pails as decribed above in 2015, and average daily water use was measured during season. Growth media was watered to field capacity and daily water use was measured through volumetric water content using a Campbell Scientific Hydrosense II. The median water depletion was 314 mL/day and this was used as an estimate of average daily crop ET under optimal soil moisture. Irrigation amounts were as follows: 100% ET: 314 mL/day, 80% ET: 253 mL/day, 60% ET: 189 mL/day.

### Data collection & analysis

Leaves were counted weekly after seedling emergence until harvest. At harvest (69 and 80 days after planting in 2018 and 2019 respectively), leaves were separated from the roots, and roots were washed to remove potting media. Leaves were counted, and total leaf area per plant was measured using a LI-3100C leaf area meter (LI-COR Inc., Lincoln, Nebraska, USA). Leaves were then dried at 60 C for 48 hours and weighed to obtain shoot biomass per plant. Root length and diameter were measured, and then roots were sliced (to speed drying) and dried for 72 hours at 60 C then weighed to obtain root biomass per plant.

Growing degree days (GDD) were calculated from daily minimum and maximum temperatures using 1.1° C as a base temperature as per Holen & Dexter, 1997; NDAWN Center, 2020 (Eq. 1)

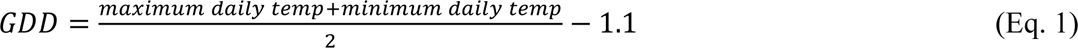

A three-parameter log-logistic model (Eq. 2) was used to model leaf development over time from emergence to harvest, where *t* is time in GDD after planting, *d* is the upper limit or maximum estimated leaves at large *t*, *e* is the inflection point of the curve, and *b* is the slope at *e*.

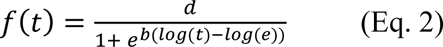

Harvest data was analyzed with a linear mixed-effects model using the lme4 package in R version 4.1.3 (Bates et al., 2015). Irrigation level, competition type, and the interaction between irrigation and competition were considered fixed effects, while year was considered a random effect. Pairwise comparisons were used to separate estimated marginal means using the emmeans and multcomp packages where appropriate (Hothorn et al., 2008; Lenth, 2022).

## RESULTS

### Shade avoidance reduced number of leaves, but not total leaf area

Sugar beet leaf production throughout the season was consistently greater without competition compared to either the shade avoidance or shade avoidance + root interaction treatments, regardless of irrigation level (Figure 2). Competition treatments explained substantially more variation in leaf numbers compared to irrigation treatments (Table 1). Averaged over irrigation treatments, leaf numbers at harvest were 23% lower in the shade avoidance + root interaction treatment compared to the no competition control (13.6 vs 17.7 leaves per plant; P < 0.001). The shade avoidance treatment (14.5 leaves per plant) had 18% fewer leaves than the no competition control (P < 0.001) when averaged over irrigation treatments. When averaged over competition treatments, leaf numbers were similar between the 100% ET and 80% ET treatments (15.7 vs 15.5 leaves per plant; P = 0.46), but the 60% ET treatment reduced leaf numbers by 6% (P = 0.013) and 7% (P = 0.002) compared to the 80% and 100% ET treatments, respectively.

**Figure 2.**
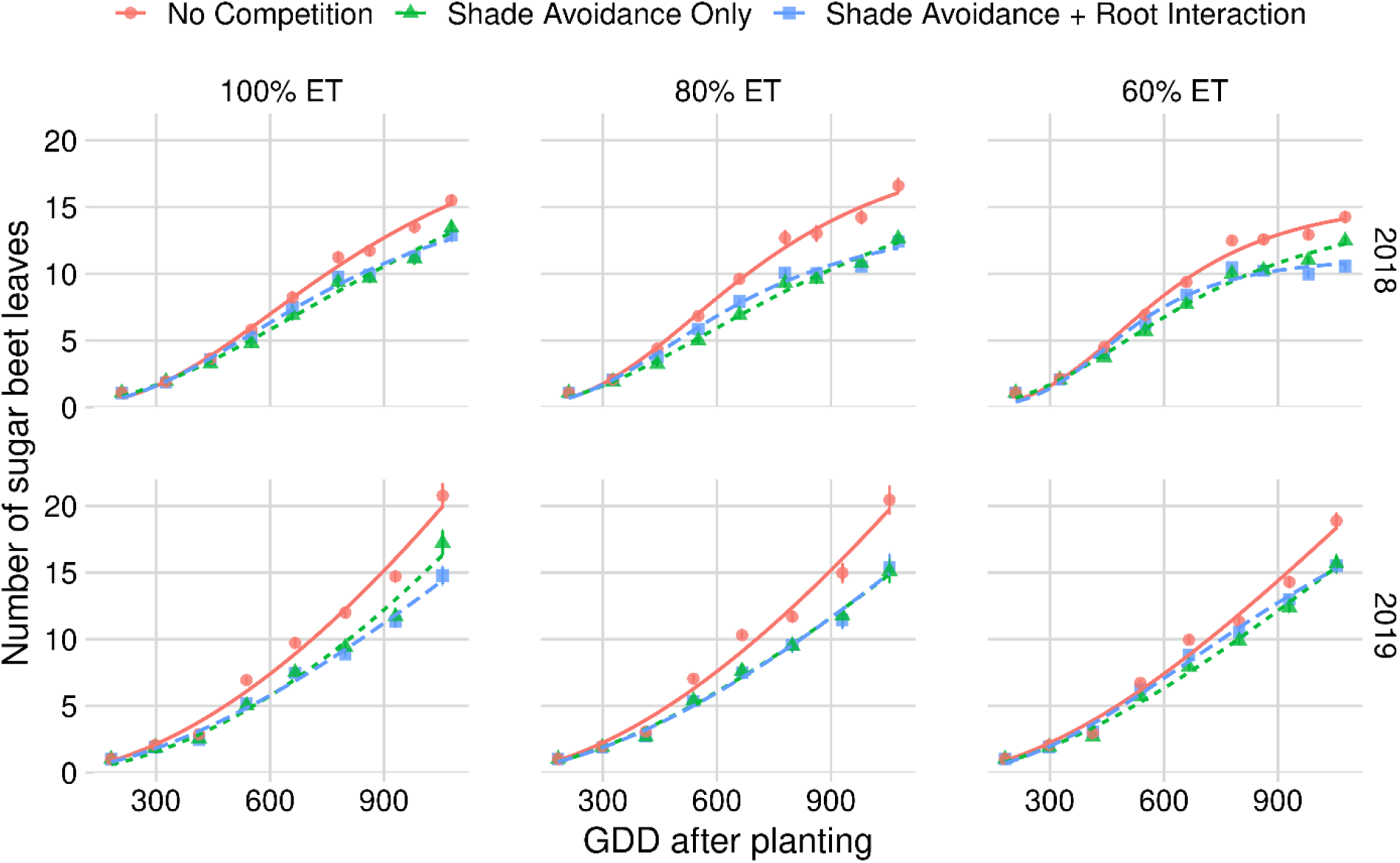
Leaf counts of sugar beets as affected by competition treatment under three different irrigation levels over two years. Vertical bars represent standard error of the mean. Model parameter estimates can be found in Supplementary Information file.

**Table 1.** Fixed effects mean squares (MS), percentage of mean squares allocated to fixed effects (%), and associated p-values for sugar beet leaf and root response variables measured at the end of the season in response to irrigation and competition treatments.

| Source of variation |  | Response variable |  |  |  |  |  |  |  |
| --- | --- | --- | --- | --- | --- | --- | --- | --- | --- |
|  |  | Leaf number | Leaf area | Leaf biomass | Root biomass | Root diameter | Root length | Specific leaf area | Root:shoot biomass |
| Irrigation | MS | 62 | 13337464 | 2196 | 2179 | 4267 | 377 | 0.296 | 2902 |
|  | % | 7% | 44% | 29% | 43% | 52% | 57% | 65% | 77% |
|  | *p-value* | 0.004 | <0.001 | <0.001 | <0.001 | <0.001 | <0.001 | <0.001 | <0.001 |
| Competition | MS | 784 | 16323022 | 5102 | 2701 | 3695 | 172 | 0.119 | 776 |
|  | % | 90% | 54% | 67% | 54% | 45% | 26% | 26% | 21% |
|  | *p-value* | <0.001 | <0.001 | <0.001 | <0.001 | <0.001 | 0.010 | 0.005 | 0.001 |
| Competition by irrigation | MS | 20 | 569656 | 345 | 167 | 244 | 107 | 0.037 | 78 |
|  | *p-value* | 0.117 | 0.083 | 0.001 | 0.039 | 0.001 | 0.022 | 0.153 | 0.545 |

There was a competition by irrigation interaction effect for sugar beet leaf area (P = 0.083), and irrigation and competition had similar magnitude effect on leaf area (Table 1). Under full irrigation, sugar beet leaf area (Figure 3b) in the shade avoidance + root interaction treatment was 31% lower than the no competition control (P < 0.001), while the shade avoidance only treatment had leaf area similar to that of the no competition control (P = 0.24). Similar effects were observed in the 80% ET irrigation treatment. However, in the 60% ET treatment, the shade avoidance treatment produced slightly *more* leaf area than the no competition control (1588 vs 1394 cm^2^, respectively; P = 0.049), even though they produced, on average, 2.5 fewer leaves per plant (Figure 3a).

**Figure 3.**
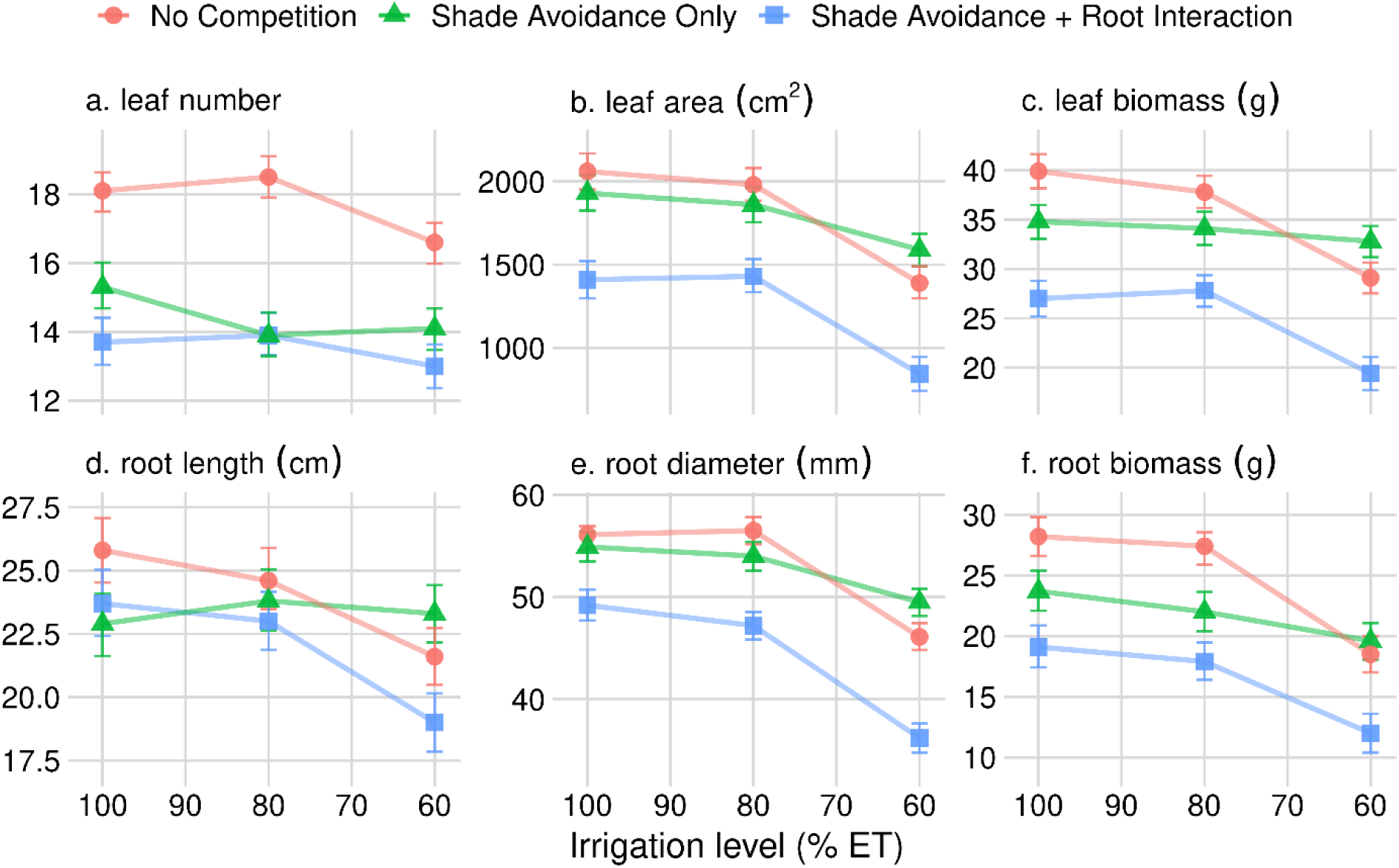
Effect of competition treatments and irrigation level on (a) sugar beet leaf number, (b) leaf area, (c) leaf biomass, (d) root length, (e) root diameter at the widest point, and (f) root biomass. Points represent estimated marginal means from linear mixed effects model. Bars are an indicator of pairwise comparisons; if bars on one point do not overlap another point, then means are statistically different (P < 0.05).

### Under severe drought stress, shade avoidance increased leaf biomass and root diameter

When fully irrigated (100% ET), the shade avoidance treatment reduced leaf biomass 13% (P = 0.004) and had similar root diameter (P = 0.42) compared to the no competition control, and results were similar under the 80% ET irrigation treatment (Figure 3c, Figure 3e). However, when irrigation was reduced to 60% of ET needs, the shade avoidance treatment produced 13% greater leaf biomass (P = 0.021) and 7% greater root diameter (P = 0.013) compared to the no competition treatment. At 80% ET, the shade avoidance treatment had 10% less leaf biomass (P = 0.029) and 20% less root biomass (P = 0.001) compared to the no competition control, but 22% greater leaf biomass (P < 0.001) and 23% greater root biomass (P = 0.011) than the root interaction treatment. Under severe drought stress (60% ET) root biomass was not significantly different among the no competition control and the shade avoidance only treatment (<1% difference, P = 0.48), even though shade avoidance reduced root biomass by 15% compared to the no competition control (P = 0.007) under 100% ET irrigation and by 19% compared to the no competition control (P = 0.001) in the 80% ET irrigation treatment (Figure 3g).

### Contrasting effects of drought stress and shade avoidance on specific leaf area and root:shoot partitioning

Specific leaf area (SLA) and root:shoot biomass allocation were influenced more by irrigation treatments compared to competition treatments (Table 1). Within each competition treatment, total sugar beet biomass (Figure 4a) and specific leaf area (SLA, Figure 4b) were reduced as drought stress increased. Averaged over irrigation treatments, the shade avoidance treatment had greater SLA compared to the no competition treatment (P = 0.005), while the combination of shade avoidance + root interaction had similar SLA as the no competition treatment (P = 0.33). When averaged over competition treatments, sugar beet root biomass allocation decreased relative to shoot allocation in the severe drought stress irrigation treatment (60% ET) compared to the 80% or 100% ET irrigation treatments. The root:shoot biomass ratios were 0.66, 0.64, and 0.58 for the 100%, 80%, and 60% ET irrigation treatments, respectively. In the no competition control, root to shoot biomass ratio increased in sugar beet at moderate drought stress (80% ET) compared to the 100% ET treatment, but then decreased substantially as irrigation was reduced to 60% ET (Figure 4c). Shade avoidance, with or without root interaction, had reduced root:shoot biomass (root:shoot = 0.61) compared to the no competition (root:shoot = 0.67; P ≤ 0.006).

**Figure 4.**
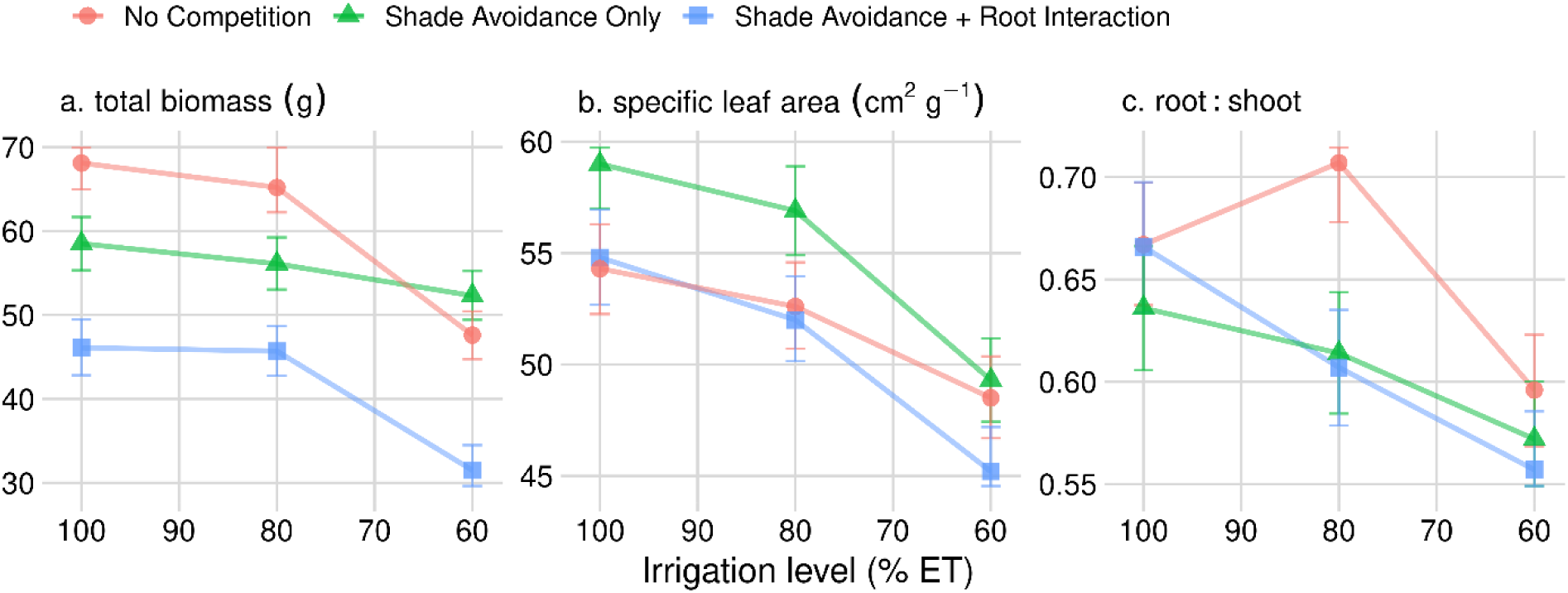
Effect of competition treatments and irrigation level on (a) total sugar beet biomass, (b) sugar beet specific leaf area, and (c) root:shoot biomass partitioning. Points represent estimated marginal means from linear mixed effects model. Bars are an indicator of pairwise comparisons; if bars on two points do not overlap, then means are statistically different (P < 0.05).

DiscussionMost previous shade avoidance research in economically important crop species have focused on tall-growing plants produced for grain, or models such as *Arabidopsis* (Casal, 2012; Green-Tracewicz et al., 2011; Page et al., 2011). Sugar beet is unique in that it is a low-growing crop selected for its resource-rich taproot, which allows us to test whether challenging the plant with opposing morphological responses would result in morphology which fits under the patterns predicted by LAM or OPT. Under no water stress, or moderate water stress, the sugar beet plant sacrifices resource stores in its taproot, but this does not happen under high water stress. Sugar beets also reduce leaf number in response to neighbor proximity but maintain leaf area and biomass mass under severe drought relative to plants grown without neighbors (Figure 2, Figure 3). To support the OPT in sugar beets, we would have expected an increase in length as predicted by previous studies. However, root length ultimately decreased in the “no competition” and “shade avoidance plus root interaction” groups, while a slight lengthening was observed in the “shade avoidance only” group at 60% ET (Figure 3d). The reduction in root length due to direct competition with other plants in the “shade avoidance plus root interaction” group was certainly driven by the lack of resources available to facilitate growth. Thus, no consistent increase in length was associated with drought growth which doesn’t necessarily fit the OPT (Figure 3d). The pattern of growth we observed was a reduction of leaf number, but no change in the leaf area (Figure 3a, Figure 3b). Overall, this translates to a reduction in structural tissue, namely petioles, while maintaining photosynthetically active tissues. Petioles are dispensable to the photosynthesis process in C3 plants like sugar beet (Johnson, 2016). Petioles are important in providing structural support to the leaves, and reducing the number of leaves could allow the sugar beet plants to devote resources into maintaining photosynthetically active parts of the leaves (Gommers et al., 2013). This reduction in structural tissue in the absence of resource competition does not seem to fit with the idea that plants allocate resources based on localized needs (Robinson, 2023).

Optimal partitioning theory (OPT) states that plants should grow in the direction of needed resources, albeit with developmental constraints (Gedroc et al., 1996; McCarthy & Enquist, 2007). This has been questioned in recent years with the advent the LAM, which states that plants depend on the resources that each organ captures locally to drive growth patterns (Robinson, 2023). Our present work challenges this latter model because plants are not constrained by resource availability under shade avoidance. Under shade avoidance, plants are only exposed to light reflected from neighboring plants. Without shading, there is no reduction in the light available to facilitate growth.

The LAM suggests that the convergence of allocation and growth-rate trajectories does not reflect an adaptive response to nutrient shortage or defoliation, but rather the influence of constraints imposed by local resource availabilities (Robinson, 2023). However, under our experimental conditions, plants undergoing the shade avoidance response without shading reduced root length and leaf numbers. This is consistent with previous findings (Adjesiwor et al., 2021; Adjesiwor & Kniss, 2020; Schambow et al., 2019). Under severe water stress (60%ET), plants reduced leaf numbers while maintaining leaf area. The LAM should predict that localized changes in resources would be driving these allocation patterns, yet we observed differences in allocation patterns without a change in localized resources.

The LAM also states that biomass allocation to roots changes when nutrients are scarce, caused by disparities in growth-rates between roots and shoots because root−shoot allocation and whole-plant growth-rate are interdependent (Robinson, 2023). In our study, as well as previous studies, light and nutrients were equally available to plants which were exposed to neighbor signals and those which were not (Adjesiwor et al., 2021; Adjesiwor & Kniss, 2020; Schambow et al., 2019). While interdependent growth patterns explain reductions in root-shoot ratio under nutrient stress, explaining altered growth patterns such as the reductions in nonessential structural tissue (e.g. petioles) without resource reduction may fall outside the predictions of the model.

The results presented here provide a good contrast to the shade avoidance responses of *Arabidopsis*, which produces smaller blades with longer petioles under shade avoidance (Casal, 2012). This is also in contrast to corn which produces smaller, but longer leaves with smaller mass (Page et al., 2009). Our expectations for this species were to allocate resources to shoot growth and our results fulfill this hypothesis, albeit in a form we did not expect. Our expectations were for growth patterns in *Beta vulgaris* to resemble those in plants such as *Arabidopsis.* While photosynthetic tissue was maintained under shade signals as expected, we did not expect to see a reduction in tissues dispensable to the photosynthesis process.

When seed of an annual species like corn is the economic product, there is a clear connection between economic yield and plant fitness because the plant’s propagules are the salable product. For sugar beet, a biennial species harvested as an annual for its root, the connection between evolutionary benefit and agronomic impact of shade avoidance is less obvious. Plant fitness is typically correlated with biomass production, and we hypothesize that the increased root biomass production under drought stress of the shade avoidance exposed plants is likely to result in greater seed production in the second year of growth (Younginger et al., 2017). Seed production in sugar beet is not an important agronomic factor, though, so it would be perhaps more interesting to evaluate this hypothesis in a different biennial species, perhaps *Beta maritima*, the progenitor species of sugar beet and other *Beta vulgaris* crop varieties like Swiss chard and table beets.

Other rosette-growing plants have been studied for their morphological responses to shade, and important differences in responses between species have been found. *Plantago lanceolata* is also a low-growing rosette and develops narrower leaves when under weed competition (Van Tienderen & Van Hinsberg, 1996). The COL-0 genotype of the model *Arabidopsis* shrinks its leaves when challenged with weedy conditions and favors the growth of petioles, which lift the blades above competing vegetation (Casal, 2012). Rhizotomatous species such as *Pachysandra* display elongated rhizomes, translating to sparse stands in shaded conditions and dense stands in open conditions (Iwabe et al., 2021). Lettuce reacts to shade avoidance by expanding its rosette, allowing more capture of light (Meng et al., 2019). In the present research, sugar beet leaves showed a decrease in total area only when challenged with moisture depletion. When taken in the context of other studies, this is a unique response and implies that the shade avoidance responses of low-growing plants is diverse and depends on a number of factors unique to each plant. We hypothesize that further studies will demonstrate that low-growing plants have diverse strategies to withstand shade which do not depend necessarily on avoiding shade by overtopping their neighbors.

Considering differences between rosette-growing plants and tall-growing plants warrants a discussion of adaptations of other plants under competitive circumstances, such as those discussed in Markestejin and Poorter (2009). While plants under drought stress are generally assumed to extend their root systems in order to obtain water from deeper soil layers, we saw no evidence of this in sugar beet taproot (Figure 3d). Plants under competitive conditions without stress are adapted to reach light by overtopping neighbors. However, sugar beets appear to respond to a combination of these conditions by growing in more resource-conservative patterns to compensate for their inability to overtop their neighbors. Thus, the generalizations made by Markestejin and Poorter (2009) may not apply to all plants, including many agronomically important species. Whether this is because sugar beet has been selected for belowground resource allocation is unknown. This hypothesis should be reexamined using plants of various and diverse growth habits. Yield loss due to weeds is a major concern to farmers, but rarely have the mechanisms of underlying losses been explored in detail (Oerke, 2006; Soltani et al., 2018). In most cases, it remains unclear what proportion of crop yield loss is driven by competitive resource use by weeds compared to light competition and attendant phenotypic responses such as shade avoidance. Phenotypic responses to light-quality cues of neighbor proximity are referred to as shade avoidance responses because, in most species, they allow plants to avoid direct shading. In a rosette-forming plant like sugar beet selected for preferential belowground allocation, the shade avoidance responses do not elevate leaf position in the canopy (Schambow et al. 2019) and therefore do not prevent the plant from being shaded. Shade avoidance provides some agronomic benefit in allowing crops to overtop and shade out competing weeds. However, higher allocation of resources to stem tissues could lead to reduced yield in some crops (Robson et al., 1996). In plants which are adapted to growing in understory habitats, shade avoidance responses tend to be selected against as excessive stem elongation is maladaptive (Dudley & Schmitt, 1996; Weinig & Delph, 2001). In this research we show that allocation adjustments arising from the perception of neighbor proximity may affect responses to concomitant environmental stresses, and, more importantly, yield at the end of the season.

One benefit of the sugar beet response to neighbor proximity becomes clear under extreme moisture limitation. Our results here suggest that ‘shade avoidance’ responses in sugar beet may actually prepare the plant for associated below-ground stressors such as water stress. Sugar beet plants maintained their ability to gather and store resources under severe drought stress relative to the no competition control, as evidenced by plants having greater leaf biomass and similar root biomass than the no competition control when exposed to shade avoidance cues (Figure 3). Reducing leaf count reduces the amount of tissue the sugar beet needs to maintain under severe drought stress, and this might be what allowed sugar beet plants to maintain root biomass, or even increase leaf mass, under shade avoidance compared to the no competition control. Although the plant’s footprint is smaller, the plants appeared able to acquire relatively more resources under severe drought stress than it would be without these shade avoidance responses. This is, therefore, not a shade *avoidance* response, as the plant is not growing tall to avoid shade. Instead, the response in sugar beet (and probably other plant species) might be better thought of more broadly as a stress tolerance response.

## ACKNOWLEDGEMENTS

Funding for this work has been provided by the USDA National Institute of Food and Agriculture (AFRI grant number 2016-67013-24912; and Research Capacity Fund WYO-631-22), and the Western Sugar Cooperative.

## AUTHOR CONTRIBUTIONS

Designed the research: ARK, BE, CW, ATA

Performed the research: ATA, JB

Collected the data: ATA, JB

Analyzed the data: ARK, JB

Wrote the manuscript: JB, ATA, ARK

Revised and edited the manuscript: ATA, CW, BE, JB, ARK

## REFERENCES

1. Adjesiwor, A. T., Ballenger, J. G., Weinig, C., Ewers, B. E., & Kniss, A. R. (2021). Plastic response to early shade avoidance cues has season-long effect on Beta vulgaris growth and development. *Plant*, Cell & Environment, n/a(n/a). 10.1111/pce.14171

2. Adjesiwor, A. T., & Kniss, A. R. (2020). Light Reflected from Different Plant Canopies Affected Beta vulgaris L. Growth and Development. Agronomy, 10(11), 1771.

3. Britz, S. J. (1990). Photoregulation of root:shoot ratio in soybean seedlings. Photochemistry and Photobiology, 52(1), 151–159. 10.1111/j.1751-1097.1990.tb01768.x

4. Casal, J. J. (2012). Shade avoidance. The Arabidopsis Book/American Society of Plant Biologists, 10.

5. Christensen, S. (1995). Weed suppression ability of spring barley varieties. Weed Research, 35(4), 241–247. 10.1111/j.1365-3180.1995.tb01786.x

6. Dudley, S. A., & Schmitt, J. (1996). Testing the adaptive plasticity hypothesis: density-dependent selection on manipulated stem length in Impatiens capensis. The American Naturalist, 147(3), 445–465.

7. Gargallo-Garriga, A., Sardans, J., Pérez-Trujillo, M., Rivas-Ubach, A., Oravec, M., Vecerova, K., Urban, O., Jentsch, A., Kreyling, J., Beierkuhnlein, C., Parella, T., & Peñuelas, J. (2014). Opposite metabolic responses of shoots and roots to drought. Scientific Reports, 4(1), 6829. 10.1038/srep06829

8. Green-Tracewicz, E., Page, E. R., & Swanton, C. J. (2011). Shade Avoidance in Soybean Reduces Branching and Increases Plant-to-Plant Variability in Biomass and Yield Per Plant. Weed Science, 59(1), 43–49. 10.1614/WS-D-10-00081.1

9. Iwabe, R., Koyama, K., & Komamura, R. (2021). Shade avoidance and light foraging of a clonal woody species, Pachysandra terminalis. Plants, 10(4), 809.

10. Jannink, J.-L., Orf, J. H., Jordan, N. R., & Shaw, R. G. (2000). Index Selection for Weed Suppressive Ability in Soybean. Crop Science, 40(4), 1087–1094. 10.2135/cropsci2000.4041087x

11. Markesteijn, L., & Poorter, L. (2009). Seedling root morphology and biomass allocation of 62 tropical tree species in relation to drought- and shade-tolerance. Journal of Ecology, 97(2), 311–325.

12. Meng, Q., Kelly, N., & Runkle, E. S. (2019). Substituting green or far-red radiation for blue radiation induces shade avoidance and promotes growth in lettuce and kale. Environmental and Experimental Botany, 162, 383–391. 10.1016/j.envexpbot.2019.03.016

13. Page, E. R., Liu, W., Cerrudo, D., Lee, E. A., & Swanton, C. J. (2011). Shade avoidance influences stress tolerance in maize. Weed Science, 59(3), 326–334.

14. Place, G. T., Reberg-Horton, S. C., Dickey, D. A., & Carter Jr., T. E. (2011). Identifying Soybean Traits of Interest for Weed Competition. Crop Science, 51(6), 2642–2654. 10.2135/cropsci2010.11.0654

15. Robson, P. R. H., McCormac, A. C., Irvine, A. S., & Smith, H. (1996). Genetic engineering of harvest index in tobacco through overexpression of a phytochrome gene. Nature Biotechnology, 14(8), 995–998.

16. Schambow, T. J., Adjesiwor, A. T., Lorent, L., & Kniss, A. R. (2019). Shade avoidance cues reduce Beta vulgaris growth. Weed Science, 67(3), 311–317. DOI: 10.1017/wsc.2019.2

17. Weinig, C., & Delph, L. F. (2001). Phenotypic plasticity early in life constrains developmental responses later. Evolution, 55(5), 930–936. 10.1111/j.0014-3820.2001.tb00610.x

18. Wille, W., Pipper, C. B., Rosenqvist, E., Andersen, S. B., & Weiner, J. (2017). Reducing shade avoidance responses in a cereal crop. AoB Plants, 9(5), plx039.

19. Younginger, B. S., Sirová, D., Cruzan, M. B., & Ballhorn, D. J. (2017). Is biomass a reliable estimate of plant fitness? Applications in Plant Sciences, 5(2), 1600094. 10.3732/apps.1600094

